# Autotrophic and mixotrophic metabolism of an anammox bacterium revealed by *in vivo* ^13^C and ^2^H metabolic network mapping

**DOI:** 10.1101/835298

**Authors:** Christopher E. Lawson, Guylaine H.L. Nuijten, Rob M. de Graaf, Tyler B. Jacobson, Martin Pabst, David. M. Stevenson, Mike S.M. Jetten, Daniel R. Noguera, Katherine D. McMahon, Daniel Amador-Noguez, Sebastian Lücker

## Abstract

Anaerobic ammonium-oxidizing (anammox) bacteria mediate a key step in the biogeochemical nitrogen cycle and have been applied worldwide for the energy-efficient removal of nitrogen from wastewater. However, outside their core energy metabolism, little is known about the metabolic networks driving anammox bacterial anabolism and mixotrophy beyond genome-based predictions. Here, we experimentally resolved the central carbon metabolism of the anammox bacterium *Candidatus* ‘Kuenenia stuttgartiensis’ using time-series ^13^C and ^2^H isotope tracing, metabolomics, and isotopically nonstationary metabolic flux analysis (INST-MFA). Our findings confirm predicted metabolic pathways used for CO_2_ fixation, central metabolism, and amino acid biosynthesis in *K. stuttgartiensis*, and reveal several instances where genomic predictions are not supported by *in vivo* metabolic fluxes. This includes the use of an oxidative tricarboxylic acid cycle, despite the genome not encoding a known citrate synthase. We also demonstrate that *K. stuttgartiensis* is able to directly assimilate extracellular formate via the Wood-Ljungdahl pathway instead of oxidizing it completely to CO_2_ followed by reassimilation. In contrast, our data suggests that *K. stuttgartiensis* is not capable of using acetate as a carbon or energy source *in situ* and that acetate oxidation occurred via the metabolic activity of a low-abundance microorganism in the bioreactor’s side population. Together, these findings provide a foundation for understanding the carbon metabolism of anammox bacteria at a systems-level and will inform future studies aimed at elucidating factors governing their function and niche differentiation in natural and engineered ecosystems.

## Introduction

Anaerobic ammonium oxidation (anammox) is a key step in the biogeochemical nitrogen cycle and represents a novel treatment process for the sustainable removal of nitrogen from wastewater. The process is mediated by a deeply branching group of chemolithoautotrophic bacteria within the Planctomycetes, the Brocadiales, that couple the anaerobic oxidation of ammonium to nitrite reduction and dinitrogen gas formation^1,2,3,4^. The discovery of this unique metabolism and subsequent translation to full-scale applications represents one of the most rapid biotechnological advances in wastewater treatment^5,6,7^. However, the metabolic networks supporting anammox metabolism remain poorly understood, which limits the prediction of their function in natural and engineered ecosystems.

Metagenomic sequencing together with experimental studies have begun to unravel the metabolic potential of anammox bacteria^2,3,8^. A combination of molecular approaches have been used to reveal the key enzymes and reactions involved in anammox catabolism, which include hydrazine (N2H4) and nitric oxide (NO) as volatile intermediates in the anammox bacterium *Candidatus* ‘Kuenenia stuttgartiensis’ (hereafter, *K. stuttgartiensis*)^3, 9, 10^. These reactions are localized within a specialized intracellular organelle, the anammoxosome, which is believed to be dedicated to energy conservation^11,12^ and also contains membrane-bound respiratory complexes of *K. stuttgartiensis*’ electron transport chain, including complex I, ATP synthase, and an NAD+:ferredoxin oxidoreductase (RNF)^13^. Experimental studies together with genomic evidence have also suggested that anammox bacteria are much more versatile than initially assumed, and can use alternative electron donors to ammonium, such as formate, acetate, and propionate for energy conservation with nitrite or nitrate as electron acceptors^2,8,14,15,16,17^. Intriguingly, it has been proposed that these organic substrates are fully oxidized to CO_2_ and not directly assimilated into cell biomass, suggesting that anammox bacteria adhere to their autotrophic lifestyle^4^.

Previous studies have shown that anammox bacteria use the Wood-Ljungdahl pathway to fix CO_2_ to acetyl-CoA based on cell carbon isotopic composition^18^, genomic evidence^2,19^ and gene expression data^4^. Based on genome annotations, four additional carboxylation reactions are also predicted to incorporate CO_2_ during central carbon metabolism, catalyzed by pyruvate:ferredoxin oxidoreductase (PFOR), 2-oxoglutarate:ferredoxin oxidoreductase (OFOR), pyruvate carboxylase, and phosphoenolpyruvate carboxylase^2,4^. Products from these reactions are proposed to flow through the tricarboxylic acid (TCA) cycle, gluconeogenesis, and the pentose phosphate pathway to synthesize all biomass precursor metabolites^2,4^. Since *K. stuttgartiensis* apparently lacks a citrate synthase gene, it has been hypothesized that the TCA cycle operates in the reductive direction via OFOR to synthesize essential precursor metabolites, such as alpha-ketoglutarate^4^. However, these genome-based predictions of *K. stuttgartiensis*’ metabolic network remain to be experimentally investigated.

Here, we resolved the central carbon metabolism of a highly enriched planktonic *K. stuttgartiensis* cell culture using time-series ^13^C and ^2^H isotope tracing, metabolomics, and isotopically nonstationary metabolic flux analysis (INST-MFA). Our results map the *in situ* central carbon metabolic network of *K. stuttgartiensis* and show that several of the genome-based network predictions summarized above are not supported by the flux of metabolites experimentally observed. For instance, *K. stuttgartiensis* operates an oxidative TCA cycle despite not encoding a canonical citrate synthase. We also demonstrate that *K. stuttgartiensis* is able to directly assimilate formate via the Wood-Ljungdahl pathway instead of oxidizing it to CO_2_ before assimilation. Moreover, our data suggests that *K. stuttgartiensis* is not capable of using acetate as a carbon or energy source *in situ* and that acetate oxidation occurred via the metabolic activity of a low-abundance microorganisms in the bioreactor’s microbial community. These findings contradict previous reports on the organic carbon metabolism of anammox bacteria and offer mechanistic insights on the versatility of carbon metabolism in *K. stuttgartiensis*.

## Results

### Mapping anammox autotrophic metabolism

To elucidate the central carbon metabolic network of *K. stuttgartiensis* under chemolithoautotrophic growth conditions, we first performed timeseries isotopic tracer experiments with ^13^C-bicarbonate coupled to metabolomic analysis. Planktonic *K. stuttgartiensis* cells were cultivated under steady-state conditions in a continuous-flow membrane bioreactor using minimal media supplemented with ammonium and nitrite. Subsequently, ^13^C-labelled bicarbonate was rapidly introduced into the bioreactor to a concentration of 30 mM (approximately 65% ^13^C-dissolved inorganic carbon, DIC), which incorporated into *K. stuttgartiensis*’ metabolome over time. Samples were collected over a 3-hour period to trace metabolic network structure based on rates of metabolite ^13^C-enrichment.

Based on the proposed carbon assimilation pathways for anammox bacteria^2,4^, we expected that CO_2_ fixation would largely occur through the Wood-Ljungdahl pathway and PFOR, resulting in fast labelling of acetyl-CoA and pyruvate, followed by phosphoenolpyruvate and other downstream metabolites. While complete labelling of phosphoenolpyruvate was observed almost immediately, ^13^C-enrichment of acetyl-CoA and pyruvate was slow during the 3-hour experiment (Figure 1a; Figure 1b). One hypothesis for this observation is substrate channeling, where the product of one enzymatic reaction is directly passed to the next enzyme without opportunity to equilibrate within the cytoplasm^20^. An alternative explanation could be that different pools of acetyl-CoA and pyruvate exist through intracellular compartmentation, which would effectively dilute the measured overall ^13^C enrichment if one pool were inactive. Consistent with the latter, amino acids synthesized from pyruvate (i.e., valine, and alanine) showed faster labelling and higher ^13^C-enrichment (Figure 1a; Figure 1b) than the pyruvate pool. This suggests that the Wood-Ljungdahl pathway and PFOR activities could occur in one compartment or specific cytoplasmic location^21^ in *K. stuttgartiensis*, while other pools of acetyl-CoA and pyruvate that do not get labelled exist in another.

**Figure 1.**
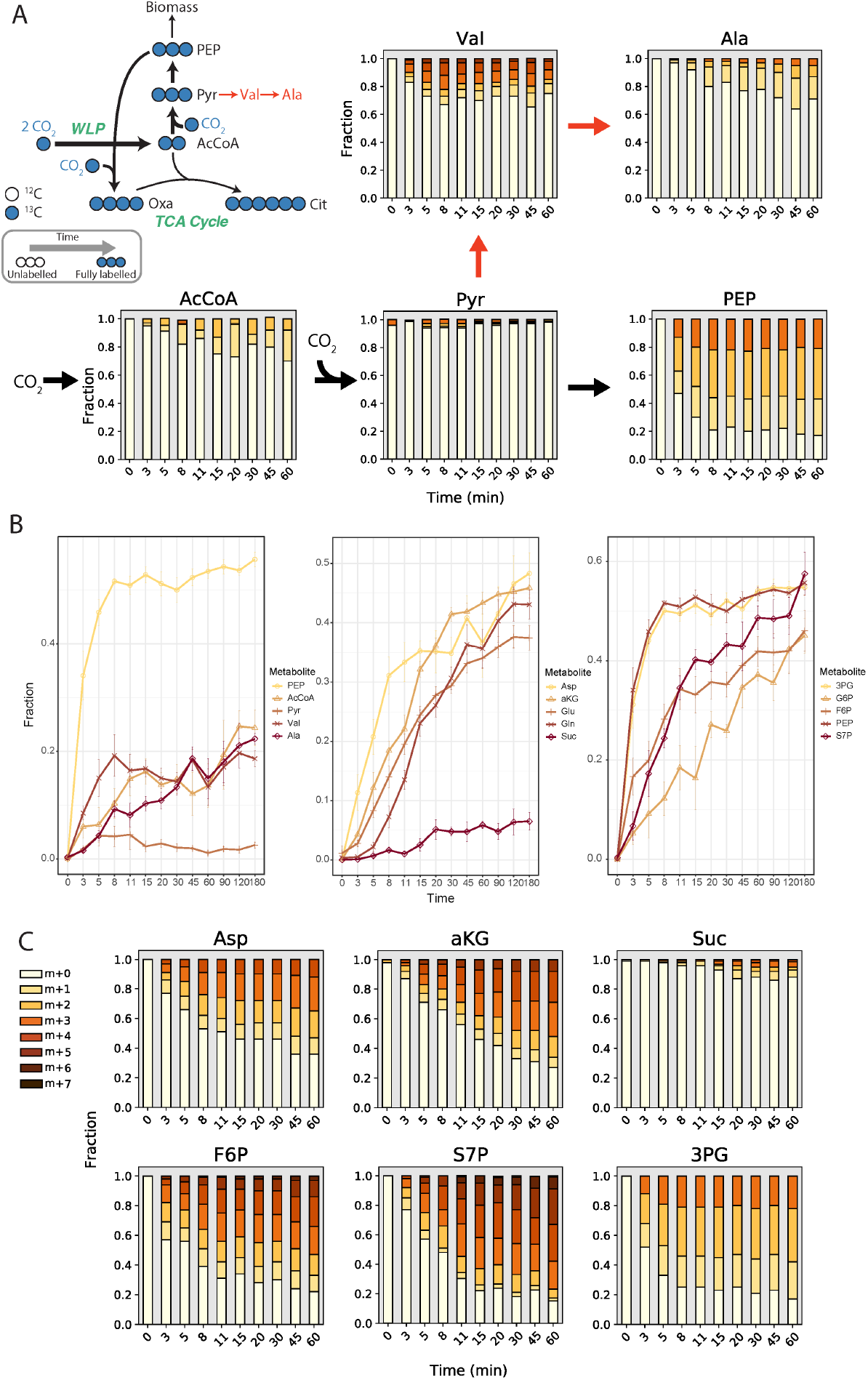
^13^C-enrichment of selected metabolites during ^13^C-bicarbonate dynamic tracing experiments. (A) Mass isotopomer distributions (MID) for selected metabolites illustrating the effects of potential substrate channeling or compartmentation of the Wood-Ljungdahl Pathway and PFOR. (B) ^13^C enrichment of metabolites associated with (left) initial CO_2_ fixation reactions (Wood-Ljungdahl Pathway, pyruvate:ferredoxin oxidoreductase) and metabolites downstream of pyruvate; (center) TCA cycle metabolites; (right) gluconeogenesis and pentose phosphate pathway metabolites. (C) Selected mass isotopomer distributions for metabolites of the TCA cycle, gluconeogenesis, and the pentose phosphate pathway. All measured metabolite MIDs represent the average of 3 independent biological replicate experiments. Metabolite MIDs and standard errors can be found in Supplementary Dataset 1.

Following CO_2_ fixation, acetyl-CoA and pyruvate are expected to enter the TCA cycle and gluconeogenesis, respectively, to produce biomass precursors. Since *K. stuttgartiensis*’ genome does not encode a canonical citrate synthase required to operate the oxidative TCA cycle, it is hypothesized that synthesis of key precursor metabolites, including succinyl-CoA and alpha-ketoglutarate, occurs via the reductive direction^4^. If this hypothesis is correct, we would expect to observe high ^13^C-labelling of oxaloacetate, succinate, and alpha-ketoglutarate. While fast labeling of aspartic acid, which was used as a proxy for oxaloacetate labelling, implied high activity of phosphoenolpyruvate (or pyruvate) carboxylase, ^13^C-labelling of succinate was much less and slower than the labelling of alpha-ketoglutarate (Figure 1b; Figure 1c). This suggested that OFOR and other reductive TCA cycle enzymes were not operating in *K. stuttgartiensis*, and that the oxidative TCA cycle was likely used to synthesize alpha-ketoglutarate.

Other biomass precursors are additionally predicted to be synthesized from gluconeogenesis and the pentose phosphate pathway in *K. stuttgartiensis*^2^. Consistent with this, we observed fast ^13^C-labeling of the gluconeogenic intermediates 3-phosphoglycerate, fructose 6-phosphate, and glucose 6-phosphate, as well as pentose phosphate pathway intermediates, such as sedheptulose 7-phosphate and ribose 5-phosphate (Figure 1b; Figure 1c; Supplementary Dataset 1).

### ^13^C-formate tracing confirms formate assimilation pathways and oxidative TCA cycle in *K. stuttgartiensis*

We further probed central carbon metabolism with ^13^C-formate. While it has been proposed that anammox bacteria fully oxidize organic substrates, such as formate, to CO_2_^4^, we hypothesized that formate could be assimilated by *K. stuttgartiensis* via the methyl branch of the Wood-Ljungdahl pathway. This would result in a positionally labelled acetyl-CoA pool that would provide additional information on metabolic network activity (Figure 2a).

**Figure 2.**
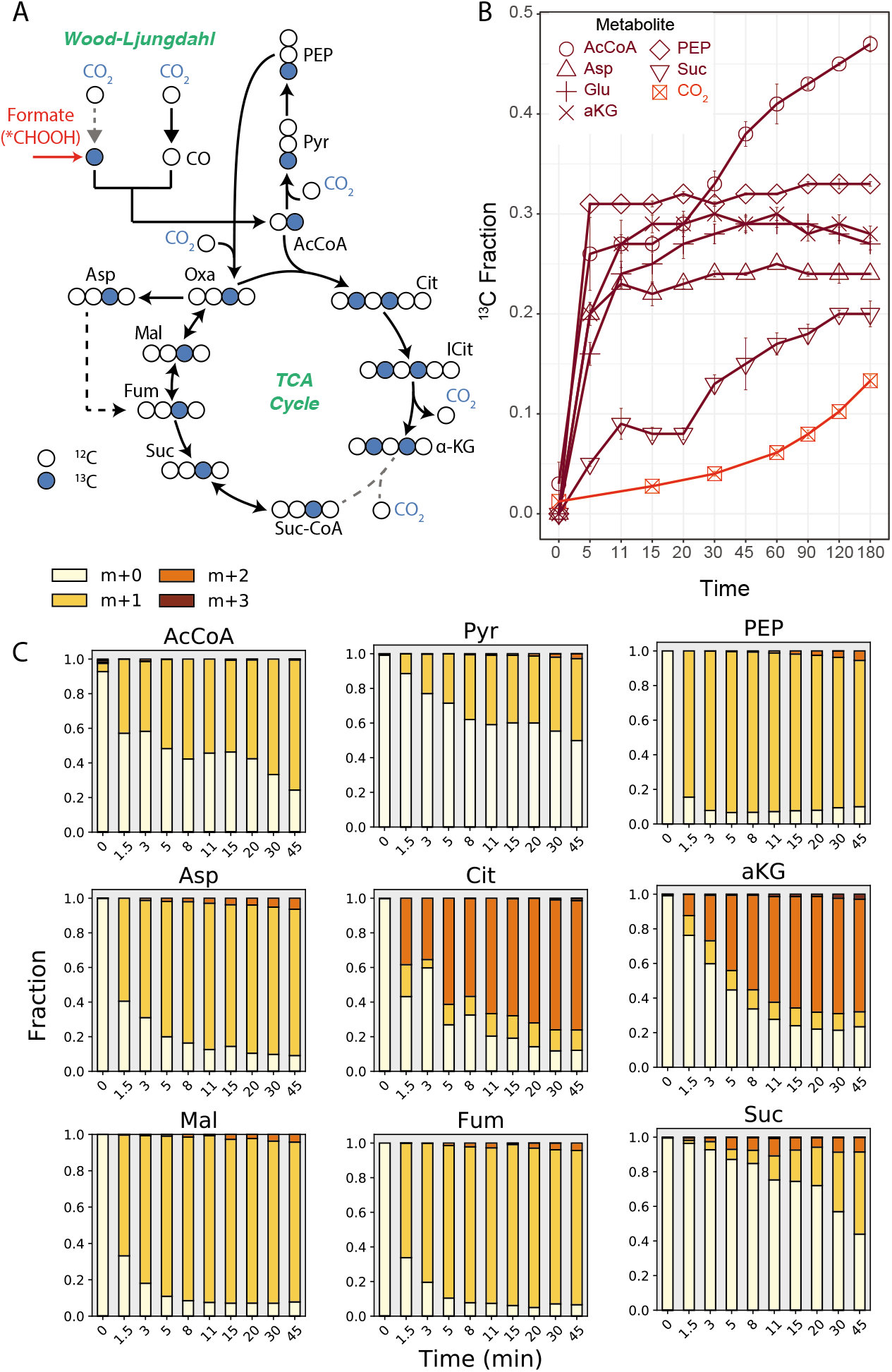
Elucidating TCA cycle of *K. stuttgartiensis* with ^13^C-formate. (A) Proposed labelling of TCA cycle metabolites with ^13^C-formate. (B) ^13^C-enrichment of selected metabolites during isotope tracer experiments with ^13^C-formate. (C) Time-series mass isotopomer distributions of selected TCA cycle metabolites during isotope tracer experiments with ^13^C-formate. All measured metabolite MIDs represent the average of 3 independent biological replicate experiments. Metabolite MIDs and standard errors can be found in Supplementary Dataset 1.

We tested this hypothesis by rapidly introducing ^13^C-formate into fresh continuous cultures of *K. stuttgartiensis* to a concentration of 50 mM followed by metabolome sampling over 180 minutes (14 timepoints total). Within 1.5 minutes of ^13^C-formate introduction, we observed steady-state labelling of several central metabolites, including phosphoenolpyruvate (Figure 2b), consistent with direct assimilation of formate. In agreement with formate assimilation via the Wood-Ljungdahl pathway, only the M+1 mass isotopomer of acetyl-CoA became enriched during the experiment (Figure 2c). M+1 mass isotopomers of phosphoenolpyruvate and aspartic acid (proxy for oxaloacetate) were also dominant (Figure 2c), consistent with their synthesis from acetyl-CoA via the sequential reactions of PFOR, phosphoenolpyruvate synthase, and phosphoenolpyruvate (or pyruvate) carboxylase, respectively. Since only a very minor fraction of the M+2 mass isotopomer were detected in these metabolites (<3% over initial 45 minutes), it can be concluded that intracellular ^13^C-CO_2_ concentrations remained low during the experiment. Consistent with this, measured ^13^C-DIC content in the liquid media produced from ^13^C-formate oxidation was low, increasing to only 5% over 45 minutes (Figure 2b). This supports the inference that ^13^C-inorganic carbon incorporation into metabolites was insignificant compared to the rate of incorporation via ^13^C-formate. Similar to ^13^C-bicarbonate experiments, slower labelling of acetyl-CoA and pyruvate was observed during the ^13^C-formate tracer experiments (Figure 2b; Figure 2c) compared to downstream metabolites. This further supports the hypothesis that separate intracellular pools of these metabolites may exist in *K. stuttgartiensis*.

^13^C-formate labelling experiments also allowed us to analyze operation of the TCA cycle. If the reductive TCA cycle was operating in *K. stuttgartiensis* only a single carbon in alpha-ketoglutarate would be labelled (from oxaloacetate), while two carbons would be labelled if alpha-ketoglutarate was produced oxidatively (from oxaloacetate and acetyl-CoA). Consistent with the latter route, mass isotopomer distributions for citrate and alpha-ketoglutarate consisted largely of M+2 mass isotopomers (Figure 2c). This clearly demonstrates that alpha-ketoglutarate was produced via an oxidative TCA cycle in *K. stuttgartiensis*, and not via the reductive branch. On the contrary, malate, fumarate, and succinate pools were largely comprised of M+1 mass isotopomers (Figure 2c), which suggests that a bifurcated TCA cycle was operating.

The labelling patterns of TCA cycle metabolites suggest that *K. stuttgartiensis* uses a novel or highly divergent enzyme for citrate synthesis. While no citrate synthase is annotated in the *K. stuttgartiensis* genome, several acyltransferase candidates exist, including genes annotated as (R)-citramalate synthase (KSMBR1_RS19040) believed to be involved in isoleucine biosynthesis^22^ and redundant copies of 2-isopropylmalate synthase (KSMBR1_RS18315 and KSMBR1_RS10820). In particular, one of the 2-isopropylmalate synthase genes (KSMBR1_RS10820) is phylogenetically related (55.1% identity) to the *Re*-citrate synthase identified in *Clostridium kluyveri*^23^ and is located next to an ADP-forming succinate-CoA ligase of the oxidative TCA cycle. Therefore, we posit that this gene encodes a dedicated *Re*-citrate synthase that allows the oxidative TCA cycle to be operational in *K. stuttgartiensis*.

### Amino acid biosynthetic pathways

^13^C-formate tracer results were also used to confirm major amino acid biosynthetic pathways in *K. stuttgartiensis*, which were largely consistent with the genome-derived predictions. Our data supports the synthesis of aspartate, asparagine, and threonine via canonical routes from oxaloacetate; the synthesis of glutamate glutamine, proline, and arginine from alpha-ketogluturate; and the synthesis of serine from 3-phosphoglycerate (Supplementary Figure 1). Furthermore, labelling patterns supported the production of valine, alanine, and leucine via canonical branched chain amino acid biosynthesis pathways from pyruvate, and of the aromatic amino acids phenylalanine and tyrosine from erythrose 4-phosphate and phosphoenolpyruvate via the shikimate pathway (Supplementary Figure 1). Interestingly,isoleucine biosynthesis was not supported by canonical routes from threonine, but rather via a recently described citramalate-dependent pathway from acetyl-CoA and pyruvate^24,25^ (Supplementary Figure 1).

Despite the *K. stuttgartiensis* genome lacking an annotated pathway for methionine biosynthesis, methionine was labelled during both ^13^C-formate and ^13^C-bicarbonate experiments. Canonical precursors for methionine biosynthesis include aspartic acid and methyl-THF (from formate via the methyl-branch of Wood Ljungdahl pathway), thus if this pathway were operating, we would expect to see mainly M+2 methionine. However, a considerable pool of M+1 methionine was consistently observed in our experiments (Supplementary Figure 1), suggesting that a potentially novel pathway to synthesize methionine is operating in *K. stuttgartiensis*, which remains to be elucidated.

### Isotopically non-stationary metabolic flux analysis of autotrophic growth

To quantitatively examine *K. stuttgartiensis*’ central carbon metabolism and obtain intracellular flux measurements, we performed INST-MFA by fitting measured, time-resolved metabolite mass isotopomer distributions from ^13^C-formate tracer experiments to an isotopomer network model^26^. This provided a quantitative systems-level flux map of *K. stuttgartiensis*’ inferred central carbon metabolism (Figure 3). Flux values were normalized to a net CO_2_ uptake rate that was estimated from the growth rate and cell carbon content: 0.0042 hrs^-1^ x 45 mmol-C/gDW = 0.186 mmol-C/gDW/hr. The resulting flux map reproduces the high intracellular flux anticipated through the Wood-Ljungdahl pathway, PFOR, and phosphoenolpyruvate (or pyruvate) carboxylase, which are the main CO_2_ fixation reactions that we observed in *K. stuttgartiensis* (Figure 3). INST-MFA also supported alpha-ketoglutarate production via the oxidative TCA cycle. Moreover, instead of running a bifurcated TCA cycle, the INST-MFA analysis predicts that the M+1 isotopomers of fumarate, succinate, and malate were indirectly derived from aspartic acid as a result of histidine and arginine biosynthesis (Figure 3). This suggests that the TCA cycle in *K. stuttgartiensis* operates incompletely, essentially functioning to produce alpha-ketoglutarate (amino acid precursor) and recycle intermediates of amino acid biosynthesis. Considerable oxidative pentose phosphate pathway flux was also measured (Figure 3). As no transhydrogenase could be identified in the genome, it is likely that this pathway is key for NAPDH generation in *K. stuttgartiensis*.

**Figure 3.**
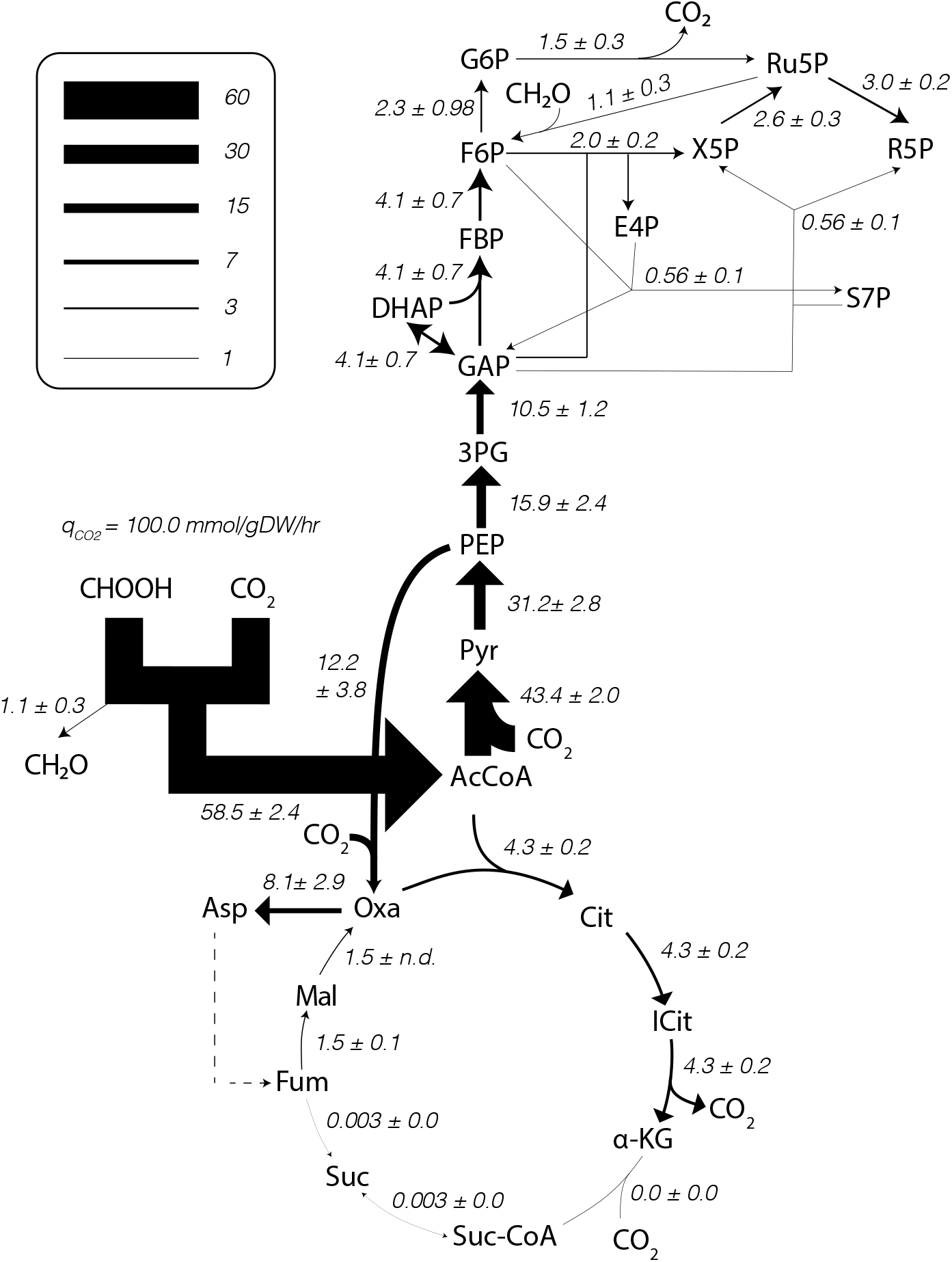
*K. stuttgartiensis* flux map generated by ^13^C INST-MFA. *K. stuttgartiensis* flux map under anaerobic, continuous flow, ammonium and nitrite medium conditions determined by fitting metabolites labelled with ^13^C-formate tracers to a single, statistically acceptable isotopomer network model. Flux values represent the net flux through a given reaction +/- standard error defined at 95% confidence. All fluxes are normalized to a net CO_2_ uptake rate of q=100 mmol-C/gDW/hr (actual CO_2_ uptake rate was 0.186 mmol-C/gDW/hr). All isotopomer network model reactions are provided in Supplementary Dataset 2. INST-MFA solutions are provided in Supplementary Dataset 3.

INST-MFA also allowed us to further resolve metabolic pathways used for the biosynthesis of sugar phosphates in *K. stuttgartiensis*. Results from the ^13^C-formate tracer experiments revealed that a considerable fraction of fructose 6-phosphate was present as M+1 mass isotopomers, which was unexpected as gluconeogenesis would result in largely M+2 mass isotopomers (Supplementary Figure 2). This suggested that alternative pathways may be operating produce fructose 6-phosphate. The genome annotation of *K. stuttgartiensis* has genes encoding for hexulose 6-phosphate synthase and 6-phospho-3-hexuloisomerase (KSMBR1_RS05220 and KSMBR1_RS18790, respectively), which are key enzymes of the ribulose monophosphate (RuMP) pathway, a formaldehyde assimilation pathway in many methylotrophic bacteria^27^. Together, these reactions fix formaldehyde to fructose 6-phosphate via a hexulose 6-phosphate intermediate^27^. We hypothesize that these reactions, as well as an unidentified formaldehyde dehydrogenase, could explain the considerable M+1 pentose and hexose phosphate isotopomers observed during ^13^C-formate labelling (Supplementary Figure 3). Consistently, including these reactions in our INST-MFA improved the model’s fit by approximately 15% (SSR of 839.9 versus 988.7, 95% confidence interval), and the model predicted that they accounted for approximately 23% of the flux synthesizing fructose 6-phosphate (Figure 3).

### ^13^C and ^2^H acetate tracing suggests *K. stuttgartiensis* does not utilize acetate as a substrate *in situ*

In addition to formate, we also examined the impact of acetate on *K. stuttgartiensis*’ metabolic network. While it has been proposed that anammox bacteria can oxidize acetate to CO_2_^8,28^, the pathways used for acetate oxidation and whether or not acetate is assimilated into biomass have yet to be resolved. If acetate were oxidized to CO_2_, we expected that it would initially be incorporated into acetyl-CoA based on previous enzymatic studies with AMP-forming acetyl-CoA synthetase (KSMBR1_RS14485)^29^, followed by oxidation to CO_2_ via either the oxidative TCA cycle or reversal of the Wood-Ljungdahl pathway, as previously suggested for other anaerobic chemolithoautotrophic bacteria^30,31^ (Figure 4a).

**Figure 4.**
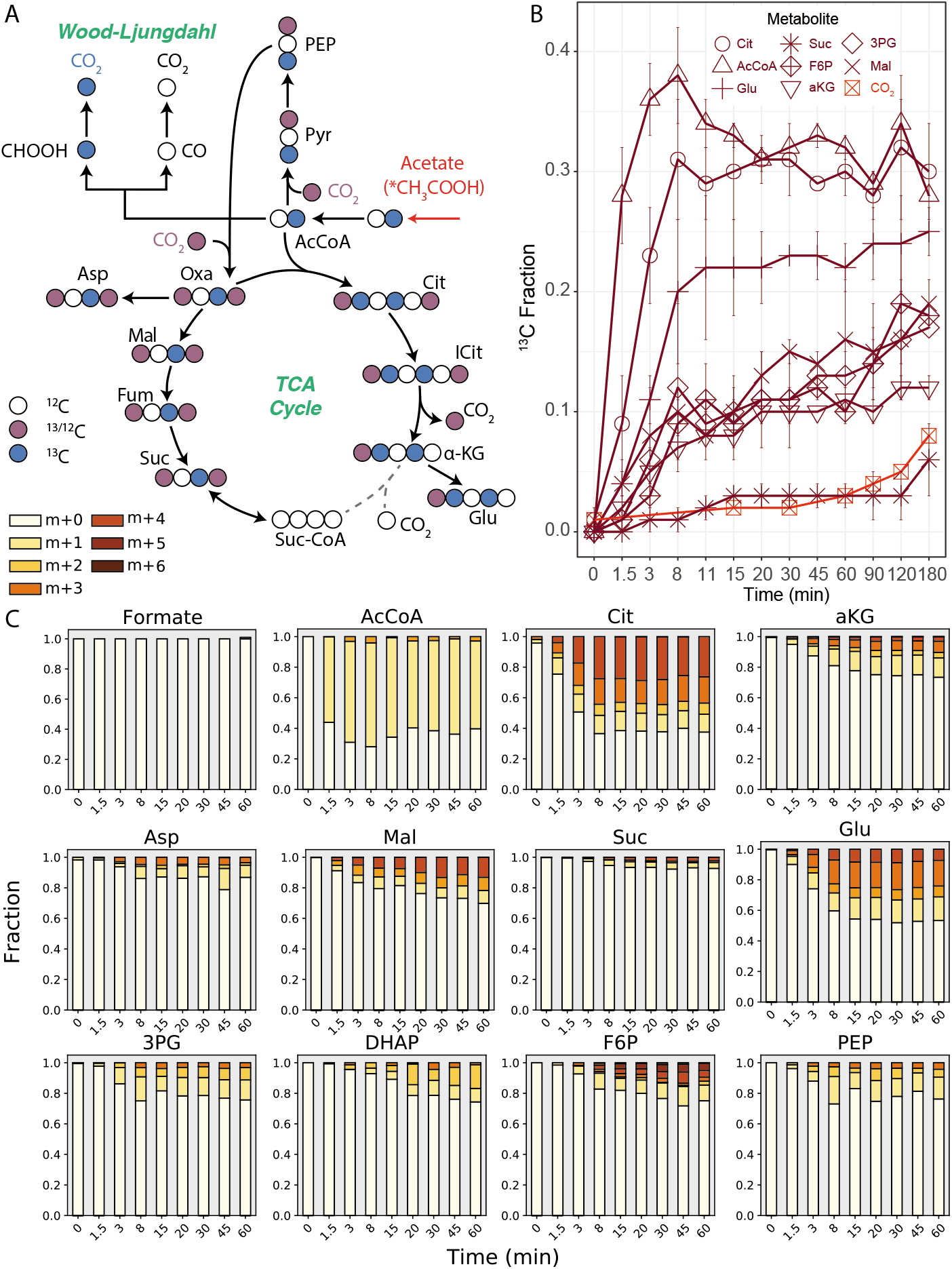
^13^C-acetate tracing reveals acetate oxidized by TCA cycle and not reverse Wood-Ljungdahl pathway. (A) Proposed labelling of TCA cycle metabolites with [2-^13^C-acetate. (B) ^13^C-enrichment of selected metabolites during isotope tracer experiments with [2-^13^C]acetate (red). (C) Time-series mass isotopomer distributions of selected TCA cycle metabolites during isotope tracer experiments with ^13^C-acetate. All measured metabolite MIDs represent the average of 2 independent biological replicate experiments, except formate (*n=1*). Metabolite MIDs and standard errors can be found in Supplementary Dataset 1.

To elucidate metabolic pathways involved in acetate metabolism, we first rapidly introduced [2-^13^C]acetate into active continuous cultures of *K. stuttgartiensis* to a final concentration of 10 mM and sampled the metabolome over 180 minutes (12 timepoints). Within 1.5 minutes after [2-^13^C]acetate addition, we observed steady-state enrichment of the M+1 mass isotopomer of acetyl-CoA, indicating its synthesis via CoA acetylation (Figure 4c). ^13^C-labelling of TCA cycle and gluconeogenic metabolites also occurred (Figure 4c), although to a much lesser extent compared to results with ^13^C-formate (Figure 2). Moreover, the mass isotopomer distributions for these metabolites appeared more evenly distributed, an observation consistent with the labeling patterns observed during ^13^C-bicarbonate tracing (Figure 1c) versus ^13^C-formate tracing (Figure 2c). This suggests that reincorporation of ^13^C-CO_2_ into central metabolism from ^13^C-acetate oxidation had occurred.

The oxidation of acetate to CO_2_ could be mediated by reversal of the Wood Ljungdahl pathway in *K. stuttgartiensis* or via the oxidative TCA cycle in either *K. stuttgartiensis* or a different organism present at low abundance in the bioreactor’s microbial community. If reversal of the Wood Ljungdahl pathway in *K. stuttgartiensis* was the main route for ^13^C-acetate oxidation, we would expect to observe an increase in ^13^C-labelled formate over time (Figure 4a). However, formate remained unlabeled during the ^13^C-acetate tracing experiments (Figure 4c), indicating the reverse Wood Ljungdahl pathway was not active and that acetate oxidation likely occurred via the oxidative TCA cycle.

To confirm this, we performed deuterium (^2^H) tracing experiments with 10 mM sodium acetate-d3, which we expected would provide clearer signal on TCA cycle activity because re-incorporation of ^13^C-CO_2_ into the metabolome would be avoided. Following ^2^H-acetate addition to the reactor, formate remained unlabeled over the course of the experiment, confirming that the reverse Wood Ljungdahl pathway was inactive. In support of acetate oxidation via the TCA cycle, ^2^H labelling of TCA cycle metabolites was observed over 180 minutes, albeit at a very slow rate (Figure 5). Interestingly, the ^2^H labelling patterns of TCA cycle metabolites were consistent with the activity of *Re*-citrate synthase, based on the presence of m+2 succinate and alpha-ketoglutarate (Figure 5), in contrast to the *Re*-citrate synthase gene candidate found in the *K. stuttgartiensis* genome. This, together with the slow increase in ^2^H labelling of TCA cycle metabolites, suggests that acetate was being oxidized by an unidentified microorganism in the bioreactor’s side population and not by *K. stuttgartiensis*.

**Figure 5.**
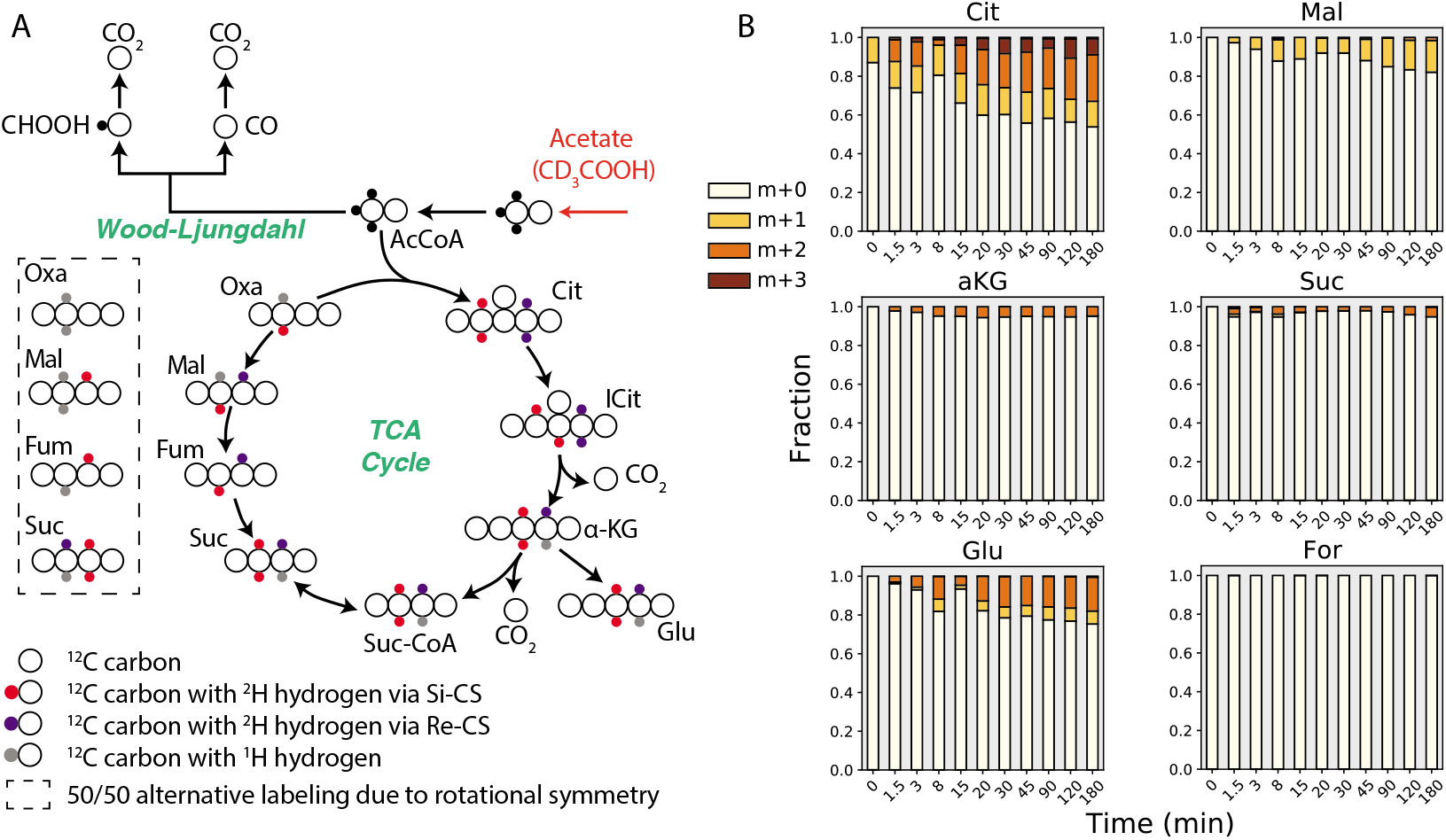
^2^H-acetate tracing suggests acetate oxidation by low abundance organism via *Si*-citrate synthase. (A) Proposed pathways and deuterium labelling for acetate oxidation via the reverse Wood-Ljungdahl pathway and TCA cycle. Black circles indicate carbons with heavy (^2^H) hydrogen, purple circles indicate carbons with ^2^H hydrogen produced via *Re*-citrate synthase, red circles indicate carbons with ^2^H hydrogen produced via. *Si*-citrate synthase. (B) Time-series mass isotopomer distributions of selected TCA cycle metabolites during isotope tracer experiments with sodium acetate-d3. All measured metabolite MIDs represent the average of 2 independent biological replicate experiments. Metabolite MIDs and standard errors can be found in Supplementary Dataset 1.

### ^13^C protein stable isotope probing confirms substrate uptake by *K. stuttgartiensis*

To confirm uptake of labelled substrates into the biomass of *K. stuttgartiensis* cells, we performed shotgun proteomics on peptides extracted from bioreactor cell pellets collected during ^13^C-labelling experiments. Metaproteomic analysis of samples collected after 0 and 72 hours confirmed that ^13^C-bicarbonate was incorporated into the *K. stuttgartiensis* proteome, increasing at a median relative isotope abundance of ~50%, consistent with the ^13^C-DIC content of the liquid media (Supplementary Figure 4). Incorporation of ^13^C-formate and [2-^13^C]acetate into the *K. stuttgartiensis* proteome was also detected after 72 hours at median relative isotope abundances of ~30% and ~10%, respectively (Supplementary Figure 4). These values are consistent with the use of the Wood-Ljungdahl pathway for formate assimilation (Figure 2), and with re-assimilation of ^13^C-CO_2_ produced from acetate oxidation, as the ^13^C-DIC in the liquid media held at ~11% between 5 and 72 hours (Figure 4b).

## Discussion

Elucidating the *in vivo* metabolic network of *K. stuttgartiensis* represents a major advance in understanding the central carbon metabolism of anammox bacteria. Our results provide the first measurements of metabolic flux via INST-MFA in an anammox bacterium, revealing a systems-level flux map of pathways involved in CO_2_ fixation, central metabolism, and amino acid biosynthesis in *K. stuttgartiensis*. We discovered that *K. stuttgartiensis* operates an oxidative TCA cycle likely mediated by a novel *Re*-citrate synthase. This pathway operates incompletely in *K. stuttgartiensis* to synthesize alpha-ketoglutarate, similar to other anaerobic bacteria^32^, and avoids the energetically costly use of reduced ferredoxin via the reductive TCA cycle. Furthermore, the considerable flux measured through the oxidative pentose phosphate pathway highlights an important link between carbon and energy metabolism for generating reducing equivalents (i.e. NADPH) in anammox bacteria.

Our analysis validated the use of the Wood-Ljungdahl pathway, PFOR, and phosphoenolpyruvate/pyruvate carboxylase for CO_2_ fixation in *K. stuttgartiensis*. We also provided first evidence of possible compartmentalization and/or metabolic channeling of these enzymes. This may enable faster pathway kinetics, avoid degradation of unstable tetrahydrofolate-based intermediates, or limit competition between competing reactions, as has been shown with other pathways^33,34^. However, further experimental investigations are required to elucidate their spatial organization in anammox cells.

In addition to CO_2_ fixation, our analysis demonstrated that *K. stuttgartiensis* can also directly use formate as a carbon source via assimilation through the Wood-Ljungdahl pathway. As formate is a common fermentation end-product found in anaerobic environments^40,41^, it is likely that anammox bacteria assimilate formate when available to lower the amount of reducing equivalents needed for CO_2_ fixation. Aside from formate, acetate has also been reported to be an organic substrate used by anammox bacteria^15,16,29,35^. However, we found no evidence that *K. stuttgartiensis* was able to use acetate as an electron donor or carbon source *in situ*, and alternatively suggest that the observed acetate oxidation activity occurred via a low-abundance heterotrophic organism present in the bioreactor’s side population. This highlights the need to further understand the metabolism of organisms that co-occur with anammox bacteria in natural and engineered ecosystems^36,37^, and to explore pathways used by other anammox species, such as Brocadia spp., which have also been reported to use acetate as a substrate *in situ*^16,28,38^.

While further studies are required to unravel the complete versatility of anammox bacteria, our study provides a foundation for understanding their carbon metabolism at a systems-level. We expect that further mapping of carbon metabolism across different anammox species and under different environmental conditions will reveal key features underlying their niche differentiation in natural and engineered ecosystems.

## Materials and Methods

### Cultivation of K. stuttgartiensis cells

A high enrichment of planktonic *K. stuttgartiensis* cells was cultivated in a continuous flow membrane bioreactor (MBR) on mineral salts medium^39^ supplemented with 45 mM of both ammonium and nitrite. The enrichment culture was estimated to be approximately 90% *K. stuttgartiensis* based on genome-centric metagenomic analysis (see Supplementary Methods). Cultures were maintained under steady-state conditions at an OD_600_ of 1.0-1.1 via continuous biomass removal and the bioreactor was continuously sparged with Ar/CO_2_ (95%/5% v/v) at a rate of 10 ml/min to maintain anaerobic conditions. The reactor hydraulic and solids retention times were approximately 46 hours and 10.5 days, respectively. The temperature and pH of the reactor were controlled at 30°C and 7.3 using a heat exchanger and 1 M KHCO_3_ buffer, respectively. The reactor was continuously stirred at 600 rpm. Nitrite concentrations were checked daily to ensure nitrite-limited conditions (Nitrite test strips MQuant^®^, Merck, Darmstadt, Germany).

### Isotope tracer experiments

Isotope tracing experiments with ^13^C- and ^2^H-labelled substrates ([^13^C]sodium bicarbonate, [^13^C]sodium formate, [2-^13^C]sodium acetate, and sodium acetate-d3; Cambridge Isotopes Laboratories, MA, USA) were performed separately on continuous cultures of *K. stuttgartiensis* cells harvested from the MBR system. ^13^C bicarbonate and ^13^C formate tracing experiments were performed in triplicate; ^13^C and ^2^H acetate tracing experiments were performed in duplicate. For each isotope tracing experiment, *K. stuttgartiensis* cells were transferred anaerobically to adjacent 1L MBRs within 10 minutes and immediately operated under identical steady-state conditions to those stated above. This resulted in no disruption of anammox activity, as measured by the absence of nitrite accumulation. Following 24 hours of steady-state operation, ^13^C-labelled substrates were rapidly introduced (within 1 minute) into the bioreactor containing *K. stuttgartiensis* cells growing under continuous cultivation. Initial reactor concentrations of ^13^C-bicarbonate, ^13^C-acetate, ^2^H-acetate, and ^13^C-formate were approximately 30 mM, 5 mM, 10 mM, and 50 mM respectively. Following ^13^C-label introduction, 5 ml samples were rapidly withdrawn from the reactor at timepoints 0, 1.5, 3, 5, 8, 11, 15, 20, 30, 45, 60, 90, 120, and 180 minutes. Samples were immediately filtered (Millipore 0.45 μm hydrophilic nylon filter HNWPO4700) using a vacuum pump to remove the medium, and filters were placed face down in 1.5 ml of −80°C extraction solvent (40:40:20 acetonitrile:methanol:water) for cell quenching and metabolite extraction. Samples were then centrifuged (10,000 rpm, 4°C, 5 mins) and 1 ml of cell-free supernatant was collected and stored at −80°C for metabolomic analysis. The time 0 min sample corresponded to the period directly before ^13^C-label addition. The ratio of ^13^C/^12^C DIC remained constant during the course of the experiment as determined by gas chromatography coupled with mass spectrometry (GC-MS) analysis (See method below).

### Metabolomic analysis

Samples were analysed using a high-performance HPLC–MS system consisting of a Vanquish^™^ UHPLC system (Thermo Scientific) coupled by electrospray ionization (ESI; negative polarity) to a hybrid quadrupole high-resolution mass spectrometer (Q Exactive Orbitrap, Thermo Scientific) operated in full scan mode for detection of targeted compounds based on their accurate masses. Properties of Full MS–SIM included a resolution of 140,000, AGC target of 1E6, maximum IT of 40 ms and scan range from 70 to 1,000 m/z. LC separation was achieved using an ACQUITY UPLC BEH C18 column (2.1 × 100 mm column, 1.7 μm particle size; part no. 186002352; serial no. 02623521115711, Waters). Solvent A was 97:3 water:methanol with 10 mM tributylamine (TBA) adjusted to pH 8.1–8.2 with 9 mM acetic acid. Solvent B was 100% methanol. Total run time was 25 min with the following gradient: 0 min, 5% B; 2.5 min, 5% B; 5 min, 20% B; 7.5 min, 20% B; 13 min, 55% B; 15.5 min, 95% B; 18.5 min, 95% B; 19 min, 5% B; 25 min, 5% B. Flow rate was 200 μl min^-1^. The autosampler and column temperatures were 4°C and 25°C, respectively. Mass isotopomer distributions were corrected for natural abundance using the method of Su et al (2017)^40^ and ^13^C enrichment values were calculated using the formula, where *N* is the number of carbon atoms in the metabolite and *Mi* is the fractional abundance of the *i^th^* mass isotopomer.

To improve separation and measurement sensitivity of specific central carbon metabolites and intracellular amino acids, samples were first derivatized with either aniline^41,42^ or benzyl chloroformate^43^, respectively. For aniline derivatization, samples were resuspended in 50 μl HPLC-grade water, 5 μl aniline (6M, pH 4.5), and 5 μl N-(3-dimethylaminopropyl)-N’-ethylcarbodiimide hydrochloride, EDC, (200 mg/ml). After 2 hours of incubation at room temperature, 1 μl of triethylamine was added to stop the reaction. For benzl chloroformate derivatization, samples were resuspended in 10 μl HPLC-grade water, 40 μl methanol, 5 μl of triethylamine, and 1 μl benzyl chloroformate and incubated at room temperature for 30 minutes.

### GC-MS analysis of dissolved inorganic carbon isotopic fractions

Isotopic fractions of DIC in the liquid media were measured based on a modified headspace method^44^. 3 ml of culture liquid were collected from the bioreactor with a syringe and directly filtered through a sterile 0.45 μm filter (Whatman, cellulose acetate) and 26G needle into a 120 ml bottle containing 1 ml 6 M HCl and crimp sealed with a rubber stopper. Prior to adding the liquid sample, bottles and HCl were flushed with either 100% N_2_ or Ar gas to void the headspace of background CO_2_. Samples were equilibrated with the acid in the bottles for at least 1 hour at room temperature to drive all DIC into the gas phase. 50 μl of the bottles headspace was then injected with a gas tight syringe (Hamilton) into a gas chromatograph (Agilent 6890 equipped with 6 ft Porapak Q columns) at 80°C with helium as a carrier gas at a flow rate of 24 ml/min, coupled to a mass spectrometer (Agilent 5975C MSD; Agilent, Santa Clara, CA) to determine the isotopic fractions of ^12^CO_2_ and ^13^CO_2_.

### Isotopic non-stationary metabolic flux analysis

Intracellular metabolic fluxes were estimated from the measured metabolite isotope labelling dynamics via INST-MFA using the elementary metabolite unit method^26^ implemented in the INCA software package v1.8^45^. Metabolic fluxes and pool sizes were estimated by minimizing the lack-of-fit between measured and computationally simulated metabolite mass isotopomer distributions using least-squares regression. All metabolite mass isotopomer distribution measurements and model reactions used for flux determination are provided in Supplementary Datasets 1 and 2, respectively. The biomass equation was based on experimental measurements of the amino acid composition obtained from *K. stuttgartiensis* biomass pellets (Supplementary Table 1). Pseudofluxes were added to the model for specific metabolites to account for inactive metabolite pools that did not participate in metabolism, but contributed to measured metabolite labelling patterns, similar to Ma et al. (2014)^46^. Chi-squared statistical tests were performed on resulting flux distributions to assess goodness-of-fit, and accurate 95% confidence intervals were computed for all estimated parameters by evaluating the sensitivity of the sum-of-squared residuals to parameter variations^47^.

### Amino acid composition analysis

Cultures were centrifuged (10,000 rpm, 15 mins, 4°C) to obtain cell pellets, which were subsequently freeze-dried prior to analysis. Total protein concentration was determined using the Pierce^™^ BCA Protein Assay Kit (ThermoFisher Scientific) and amino acid composition was determined according to Carnicer et al. (2009)^48^ using a Varian 920-LC high performance liquid chromatography amino acid analyzer.

### ^13^Cprotein stable isotope probing

Proteins were extracted from bioreactor cell pellets using glass bead beating (acid, washed, 0.1 mm diameter) in a suspension containing B-PER reagent (Thermo Scientific, Germany) and TEAB buffer (50 mM TEAB, 1% (w/w) NaDOC at pH 8). Following DTT reduction and alkylation using iodo acetamide (IAA), protein extracts were subject to proteolytic digestion using trypsin. Resulting peptides were solid phase extraction-purified using an Oasis HLB 96 well plate (Waters, UK), according to the manufacturer protocols. The purified peptide fraction was analysed via a one-dimensional reverse phase separation (Acclaim PepMap RSLC RP C18, 50 μm x 150 mm,2μm, 100A) coupled to a Q Exactive plus Orbitrap mass spectrometer (Thermo Scientific, Germany) operating in data-dependent acquisition mode (DDA, shotgun proteomics). The flow rate was maintained at 300 nL/min over a linear gradient from 5% to 30% over 90 minutes and finally to 75% B over 25 minutes. Solvent A was H_2_O containing 0.1% formic acid, and solvent B consisted of 80% acetonitrile in H_2_O and 0.1% formic acid. The Orbitrap was operated in DDA mode acquiring peptide signals from 350-1400 m/z, where the top 10 signals (with a charge between 2-7) were isolated at a window of 2.0 m/z and fragmented using a NCE of 30. The AGC target was set to 1E5, at a max IT of 54 ms and 17.5 K resolution. Protein identification and relative isotope abundances were determined from Tandem-MS data using PEAKS Studio X (BSI, Canada) and MetaProSIP (OpenMS, Univ Tuebingen/Berlin, Germany)^49^ integrated into the KNIME 4.0.1 analytics platform (Zurich, Switzerland), respectively. All peptide spectra were matched against a protein database generated from predicted open reading frames from the *K. stuttgartiensis* genome.

## Acknowledgements

The authors would like to acknowledge Patricia van Dam and Carol de Ram for help with metaproteomic sample preparation, Katinka van de Pas-Schoonen for help with bioreactor maintenance, Paul van der Ven and Sebastian Krosse for help with amino acid analysis, and Arjan Pol and Huub Op den Camp for helpful discussions. Funding was provided by the National Science Foundation (CBET-1435661, CBET-1803055 and MCB-1518130), the Netherlands Organization for Scientific Research (016.Vidi.189.050 and SIAM Gravitation Grant 024.002.002), the European Research Council (ERC Advanced Grant Ecomom 339880), a Wisconsin Distinguished Graduate Fellowship, a Postgraduate Scholarship-Doctoral (PGS-D) by the National Sciences and Engineering Research Council of Canada (NSERC), and the UW-Madison Office of the Vice Chancellor for Research and Graduate Education through the Microbiome Initiative.

## Contributions

C.E.L., S.L., D.A-N., K.D.M., and D.R.N. designed the study. C.E.L., G.N., R.G. performed the ^13^C tracer experiments. C.E.L., T.B.J., and D.M.S. performed the metabolomic analysis. M.P. and C.E.L. performed the metaproteomic analysis. C.E.L. wrote the manuscript. All authors provided valuable feedback and edits on the manuscript.

---

**Supplementary Figure 1.**
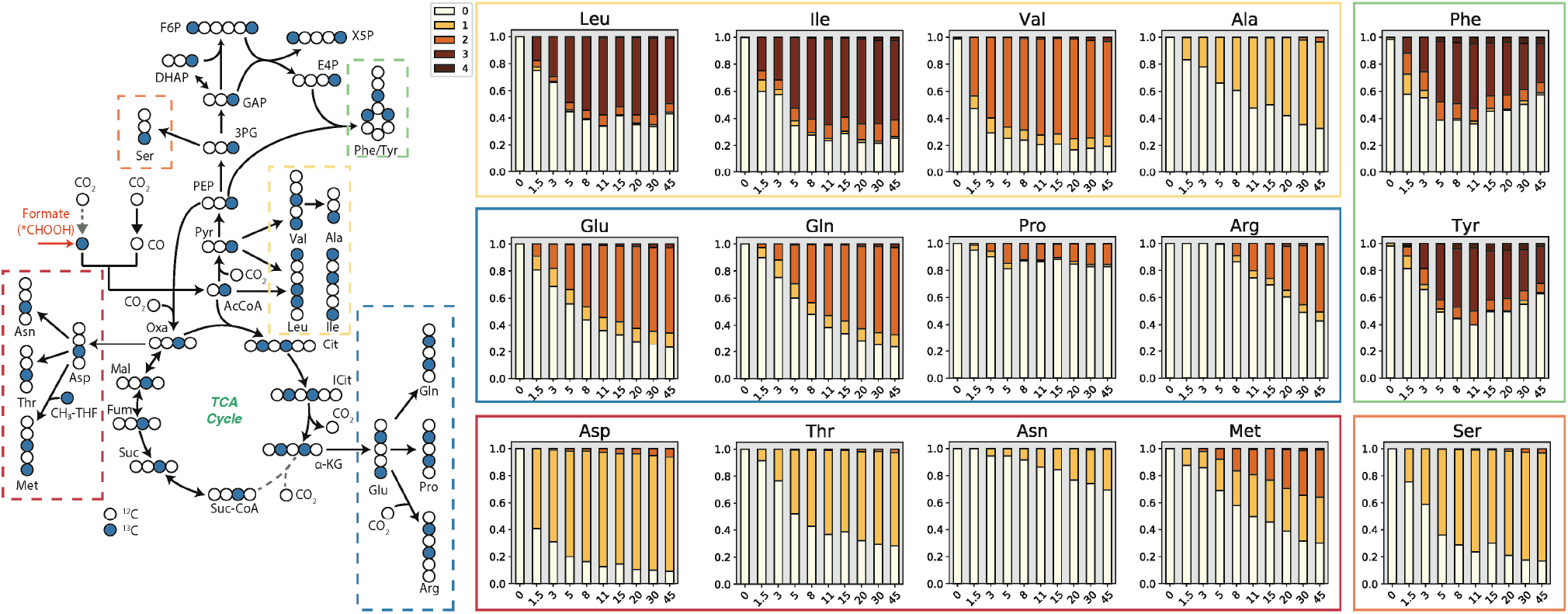
Confirmation of amino acid biosynthetic pathways in *K. stuttgartiensis*. (Left) Expected metabolite labeling patterns from ^13^C-formate. (Right) Mass isotopomer distributions for measured intracellular amino acids. All measured metabolite MIDs represent the average of 2 independent biological replicates experiments. Metabolite MIDs and standard errors can be found in Supplementary Dataset 1.

**Supplementary Figure 2.**
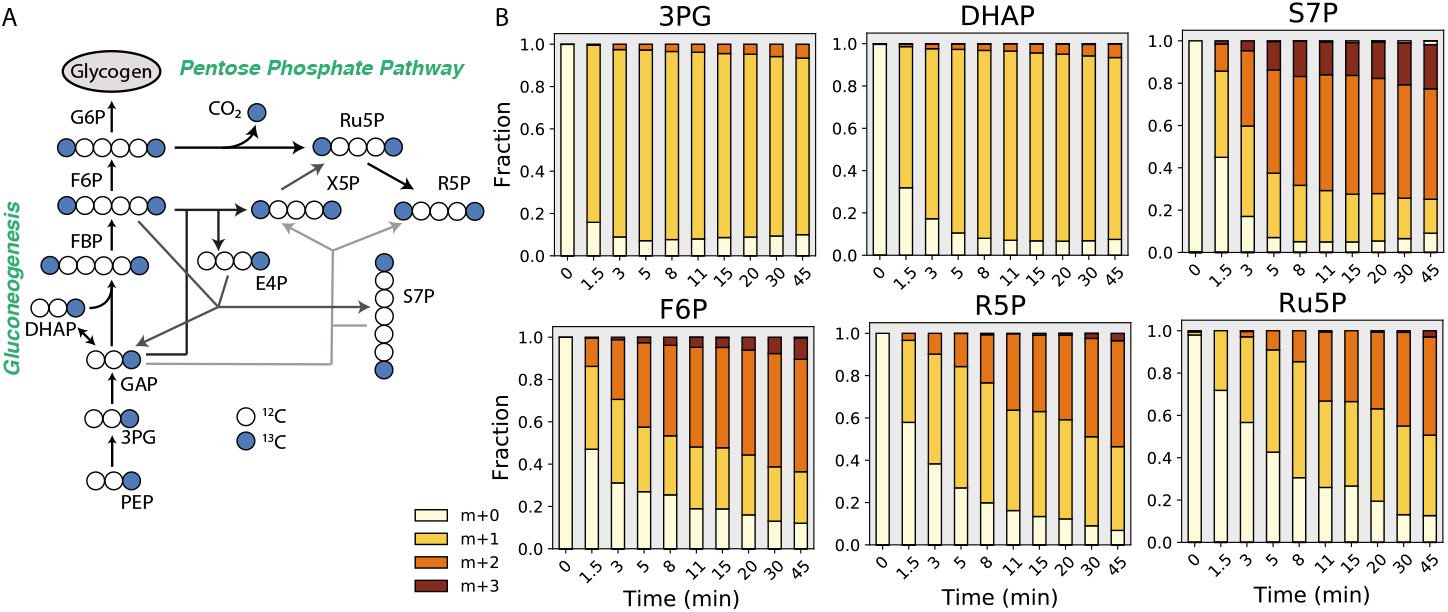
Operation of gluconeogenesis and pentose phosphate pathway in *K. stuttgartiensis* revealed by ^13^C-formate dynamic labelling experiments. (A) Proposed atom mapping of gluconeogenesis and pentose phosphate pathway from ^13^C-formate labelled phosphoenolpyruvate at steady-state. (B) Time-series mass isotopomer distributions of selected gluconeogenesis and pentose phosphate pathway metabolites during dynamic isotope tracer experiments with ^13^C-formate. All measured metabolite MIDs represent the average of 3 independent biological replicate experiments. Metabolite MIDs and standard errors can be found in Supplementary Dataset 1.

**Supplementary Figure 3.**
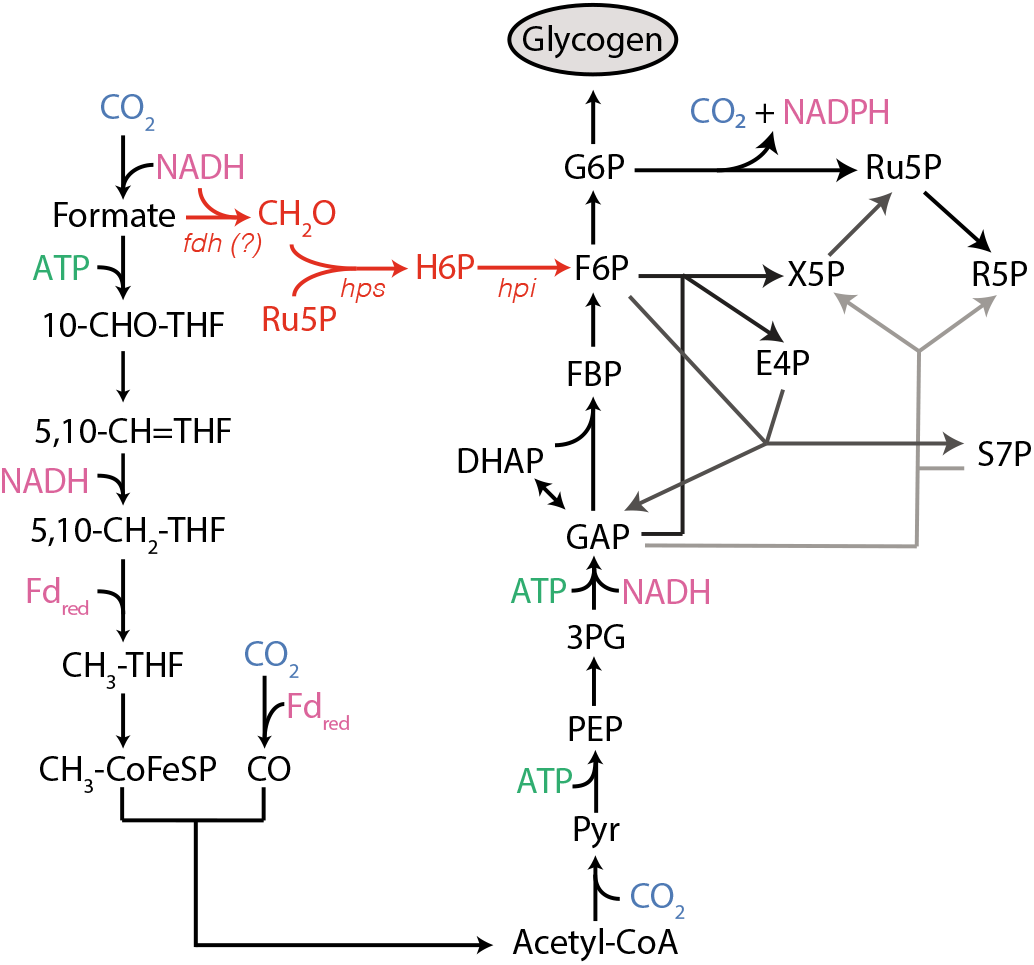
Proposed synthesis of sugar phosphates from gluconeogenesis and the RuMP cycle in *K. stuttgartiensis*. RuMP cycle reactions that synthesize fructose 6-phosphate (F6P) from formaldehyde (CH_2_O) and ribulose 5-phosphate (Ru5P) are shown in orange. Production of CH_2_O from formate via an unknown formaldehyde dehydrogenase is also show in orange. Reducing equivalents shown in pink; ATP shown in green; CO_2_ shown in blue. *fdh*: formaldehyde dehydrogenase; *hps*: hexulose 6-phosphate synthase; *hpi*: hexulose 6-phosphate isomerase; *?*: gene not identified in genome.

**Supplementary Figure 4.**
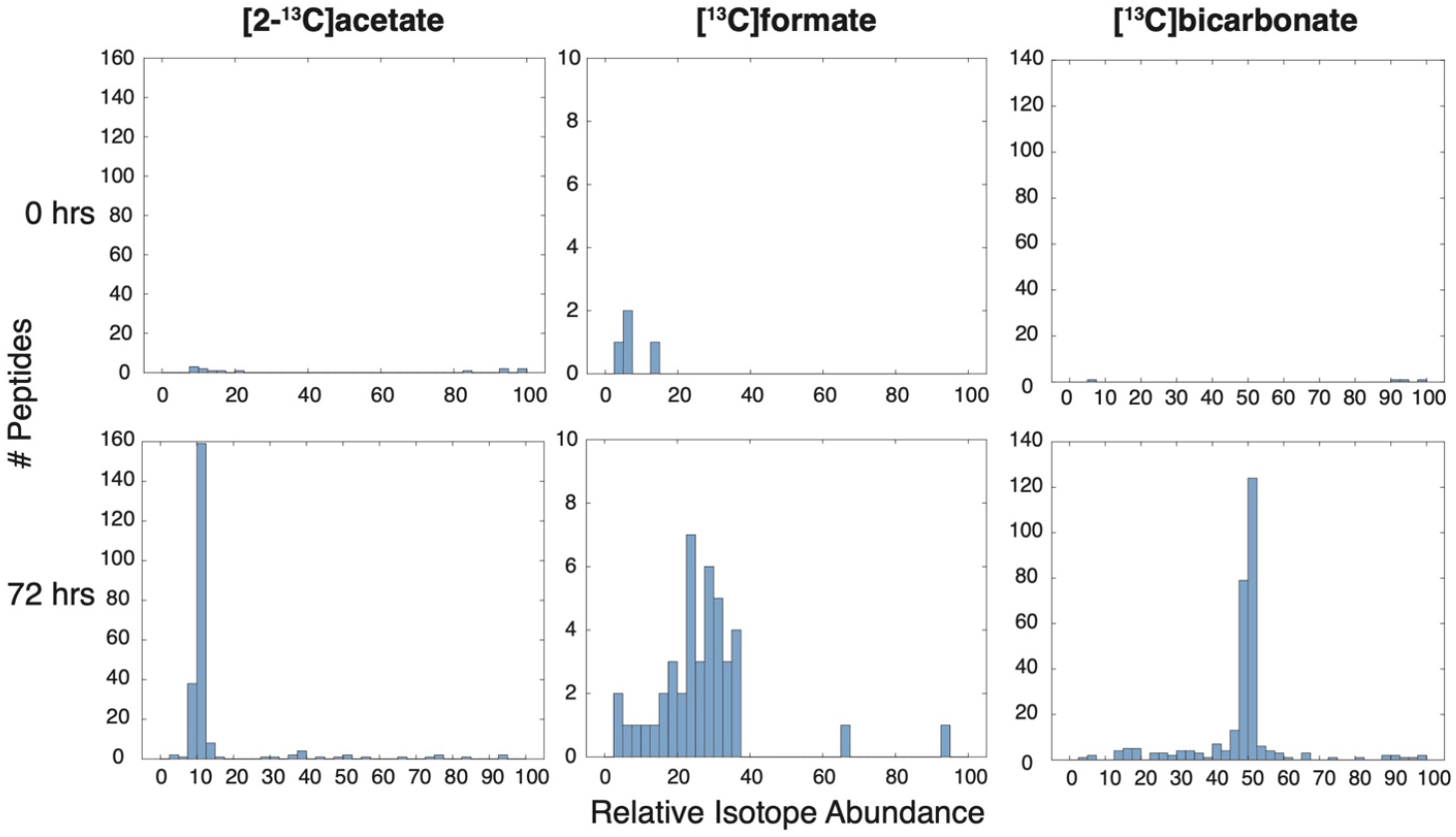
Confirmation of ^13^C-labelled substrate incorporation into the proteome of *K. stuttgartiensis*. Distribution of relative isotope abundances for identified peptides assigned to the *K. stuttgartiensis* proteome during [2-^13^C]acetate, ^13^C-formate, and ^13^C-bicarbonate tracer experiments after 0 and 72 hours.

**Supplementary Table 1.** *K. stuttgartiensis* biomass amino acid composition.

| **Amino**<br>**Acid** | **Mass**<br>**($\mu\text{mol/mgDW}$)** | **Std Dev**<br>**($\mu\text{mol/mgDW}$)** |
| --- | --- | --- |
| Ala | 374.6 | 38.7 |
| Arg | 278.9 | 56.7 |
| Asp | 130.0 | 10.2 |
| Glu | 107.5 | 8.5 |
| Gly | 778.8 | 117.4 |
| His | 42.2 | 7.6 |
| Ile | 267.8 | 43.4 |
| Leu | 393.2 | 55.3 |
| Lys | 154.8 | 44.1 |
| Met | 218.9 | 58.4 |
| Phe | 205.4 | 43.0 |
| Pro | 460.0 | 96.1 |
| Ser | 436.5 | 94.1 |
| Thr | 381.0 | 86.5 |
| Val | 438.2 | 76.2 |

**Supplementary Dataset 1.** Average metabolite mass isotopomer distributions and associated standard errors during ^13^C-bicarbonate, ^13^C-formate, and [2-^13^C]acetate tracer experiments. (Sheet 1A) Average mass isotopomer distributions for selected metabolites during 13C-bicarbonate tracing; (Sheet 1B) mass isotopomer distributions standard error values for selected metabolites during 13C-bicarbonate tracing; (Sheet 2A) average mass isotopomer distributions for selected metabolites during 13C-formate tracing; (Sheet 2B) mass isotopomer distributions standard error values for selected metabolites during 13C-formate tracing; (Sheet 3A) average mass isotopomer distributions for selected metabolites during [2-^13^C]acetate tracing; (Sheet 3B) mass isotopomer distributions standard error values for selected metabolites during [2-^13^C]acetate tracing. (Sheet 4A) Average mass isotopomer distributions for selected metabolites during sodium acetate-d3 tracing, (Sheet 4B) Mass isotopomer distributions standard error values for selected metabolites during sodium acetate-d3 tracing.

**Supplementary Dataset 2.** *K. stuttgartiensis* isotopomer network model. Letters within brackets indicate carbon atom transitions of each metabolite for a given reaction.

**Supplementary Dataset 3.** INST-MFA model results. Metabolite MIDs used for model fitting were Pro, Asn, Ala, Thr, aKG, Ser, Suc, Asp, Glu, R5P, PEP, Cit, Mal, Ru5P, Fum, F6P, Pyr, G6P, Val, CO_2_, and Gln at timepoints 0, 1.5, 3, 5, 8, 11, 15, 20, 30, and 45 minutes. All metabolite MIDs can be found in Supplementary Dataset 1.

**Supplementary Dataset 4.** Summary of the metagenome assembled genomes (MAGs) recovered from the anammox membrane bioreactor investigated in this study. Replicate 1 was collected on 18/12/2019 immediately prior to a ^13^C-formate tracer experiment; Replicate 2 was collected on 19/08/2020 immediately prior to a ^13^C-acetate tracer experiment. The recovered *K. stuttgartiensis* genome (planctomycetaceae_1_das_tool) shared 99.6% genomic average nucleotide identity to the previously published *K. stuttgartiensis* genome (NCBI ID: LT934425.1) and is highlighted in bold text.

## Notes

### Competing Interest Statement

The authors have declared no competing interest.

